# A marsh samphire (*Salicornia europaea* ) extract induces a strictly sex-specific, Tor-signaling-dependent extension of lifespan and exhibits anti-obesogenic properties

**DOI:** 10.1101/2024.08.25.609591

**Authors:** Navid Tahan Zadeh, Mirjam Knop, Lisa Maria Ulrich, Iris Bruchhaus, Roman Lang, Kai Lüersen, Gerald Rimbach, Thomas Roeder

**Author notes:** Correspondence: Prof. Dr. Thomas Roeder, Kiel University, Dept. Molecular Physiology, Kiel, Germany.

## Abstract

We have studied the health-promoting properties of the marsh samphire *Salicornia europaea* by employing the fruit fly *Drosophila melanogaster*. Supplementing a standard diet with 0.2% of an aqueous extract of this halophilic plant (SEE), which grows in sandy and muddy habitats in intertidal zones, extended the lifespan of two different *Drosophila* strains by up to a third. This effect was strictly sex-specific and only affected females. When analyzing the body composition, we found that the body fat content of SEE-treated female flies was lower compared to controls. The extract also positively impacted the lifespan of flies fed a high-fat diet but not a high-sugar diet. It is of note in this context that SEE exhibited a lipase-inhibitory activity in vitro. Moreover, SEE counteracted ageing-associated loss of intestinal barrier integrity. The sex-specific lifespan extensions induced by the SEE were entirely dependent on functional Tor signaling in the flies. Tissue-specific silencing of the Tor signalling pathway in different cellular compartments of the intestine reduced, but did not completely abolish, the lifespan-prolonging effect in females. This suggests that other organs also contribute to this effect. In contrast to the observation that Tor signalling is necessary for the life-prolonging effect, Foxo deficiency led to a reduction in the effect, but not to its complete absence. In conclusion, the SEE is a promising candidate for a health-promoting intervention, and we laid the current groundwork for identifying the compounds that mediate this effect in future studies.

## Introduction

Our diet influences essential aspects of our lives. This statement is especially true for the various facets of health. The diet’s composition, which can have positive or negative effects, is critical. Here, the research focused often on the relevance of macronutrient ratios^1–3^. In contrast, the health effects of other dietary components are far less well studied. In particular, certain secondary metabolites have already been shown to have a positive impact on human health. We can find these health-promoting nutritional components in a wide variety of sources including different plants, algae, and fungi. Especially those plants, algae and fungi whose health-promoting properties are already known from traditional use, for example, are promising sources for in-depth analyses. Usually, only a few metabolites of these natural sources are responsible for the health-promoting effects. A typical example of such a pharmacologically active metabolite is rapamycin, a macrolide isolated from the fungus *Streptomyces hygroscopicus*^4^. Rapamycin has a health-promoting and life-prolonging bioreactivity by specifically intervening in the Tor signaling pathway and thus mimicking a calorie-reduced diet^5–7^, a mechanism of action common to many lifespan-prolonging interventions.

Marine plants and algae are excellent candidates to expand the range of accessible nutritional sources that exhibit health- and life-prolonging effects. This assumption is based on the observation that healthy or particularly long-living populations have high proportions of certain marine plants and algae in their daily diet^8^. An example is the Okinawa region, where people consume a low-calorie diet with large amounts of plant-based marine products^9,10^. Building on these findings, we conducted a broad-based screening using a comprehensive plant and algal extract library ^11^ and evaluated their life-prolonging effects. We used the fruit fly *Drosophila melanogaster* as a screening model, which is optimally suited for such experimental strategies^12–14^. Thereby, three decisive advantages come together in *Drosophila*: the flies have an organ composition very similar to ours, the metabolic characteristics exhibit a high degree of similarity to ours, and it is experimentally amenable to high-throughput screening of extracts and substances in terms of their effects on longevity^15^. Comprehensive screens like these have already allowed us to identify the life-prolonging effects of extracts of the marine algae *Saccorhiza polyschides* and *Eisenia bicyclis* and to elucidate their mode of action^16–18^. In the present work, we studied the health-promoting effects of aqueous *Salicornia europaea* extracts (SEE). *S. europaea*, also known as marsh samphire, sea asparagus, glasswort, or pickleweed, is a halophytic extremophile found primarily in intertidal salt marshes, mangroves, or beaches^19^. For centuries, it was used for food and medicinal applications. It has antioxidant, anti-inflammatory, antidiabetic, and anticancer properties and can help slow aging^20,21^. It also has antibacterial activity^22^.

In this project, we studied the effects of SEE on lifespan and elucidated the underlying mechanisms by using *D. melanogaster*. We showed that SEE extends the lifespan of flies by more than 30% in a sex-specific manner only affecting females. Moreover, the extract reduced the triacylglyceride levels in flies and increased their survival on a high-fat diet, implying that the extract has lipid-lowering properties. Our data suggest that the aqueous extract of *S. europaea* contains bioactive compound(s) which target the Tor signaling pathway to mediate the lifespan extension in females.

## Results

### *Salicornia* extracts extend the lifespan of *Drosophila melanogaster* in a sex-dependent manner

We identified an aqueous extract of *S. europaea* (SEE) during a larger screen of plant and algal extracts for their life-extending potential using fruit flies as a model system. To further characterize and verify this lifespan-prolonging effect, we used two different concentrations (0.2% and 0.05%) of this extract and measured lifespan in cohorts of mated females of the *D. melanogaster w*^*1118*^ strain. Animals subjected to 0.05% SEE did not show any lifespan prolongation (Fig. 1A). However, application of the extract at the higher concentration of 0.2% significantly increased median lifespan by 36.6% (Fig. 1B, p < 0.0001). We also tested cohorts of mated males, but we did not observe any lifespan extension by supplementation with 0.2% SEE in males (Fig. 1C). To exclude that the observed effects on lifespan are only strain-specific, we tested a second *Drosophila* strain, namely *yw*. Here, we could also show a robust lifespan prolongation by 16.1% induced by 0.2% SEE (Fig. 1D, p<0.0001). The increases in lifespan were not only seen for the median lifespan but also the maximal lifespans (i.e., the 10% longest-living animals) in both *w*^*1118*^ (49 d and 53 d; p<0.0001)) and *yw* animals (43 d and 49 d, p < 0.0001).

**Figure 1:**
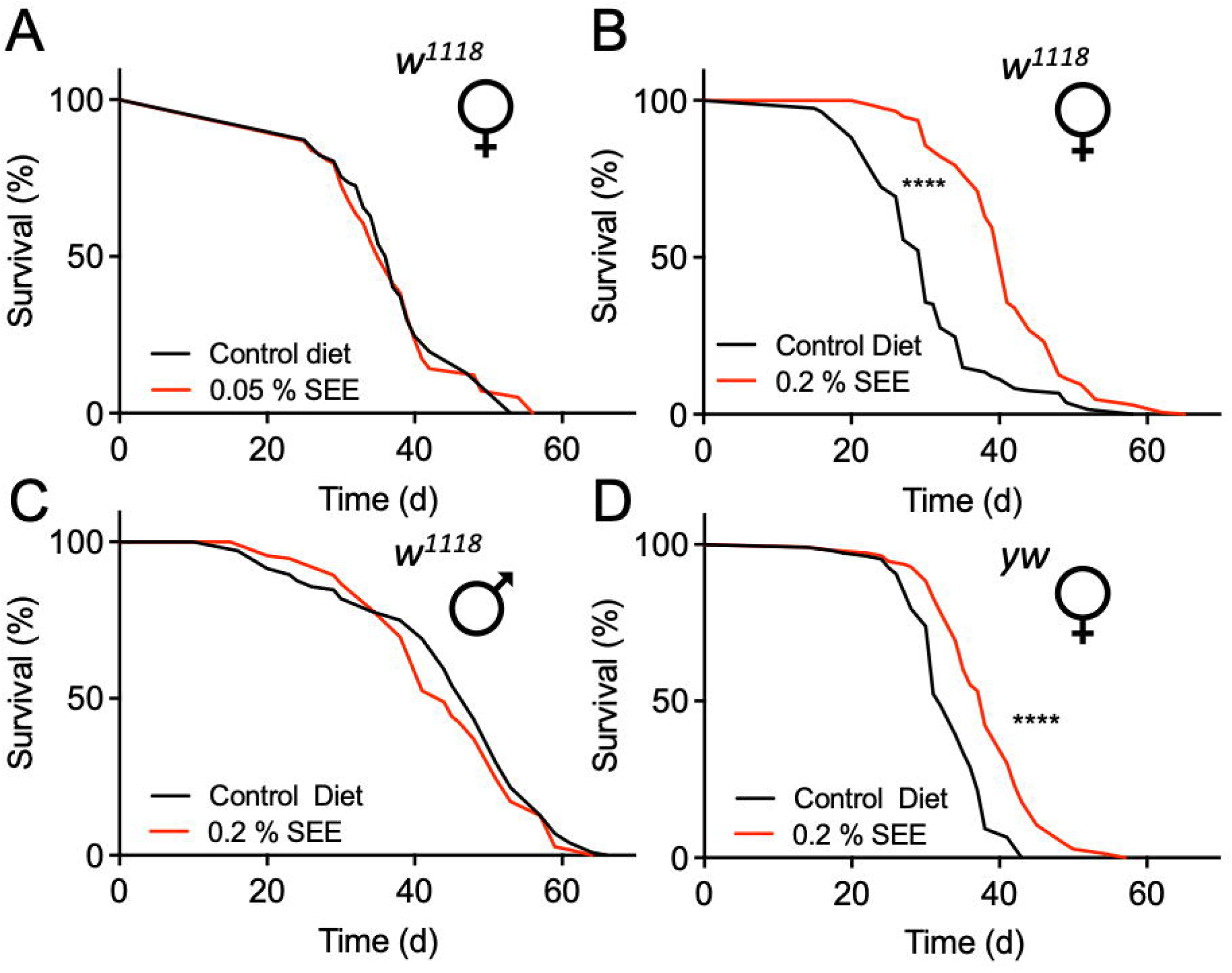
Effect of SEE on lifespan in flies of different sexes. (A) Effect of supplementation with 0.05% of SEE on lifespan of *w*^*1118*^ female flies compared with untreated animals (N = 100, the median lifespan of SEE-treated flies = 35 d, the median lifespan of untreated flies = 37 d). (B) 0.2% of SEE effects on lifespan of female *w*^*1118*^ flies (N = 150, median lifespan of SEE-treated animals = 41 d, the median lifespan of the control group = 30 d). (C) Comparison of lifespans of male *w*^*1118*^ flies subjected to 0.2% SEE and those receiving a non-supplemented diet (N = 100, median lifespan of both groups = 41 d). (D) SEE increased the lifespan of *yw* female flies (N = 100, median lifespan of flies treated with SEE = 38 d, median lifespan of control = 32 d). SEE: *Salicornia europaea* extract, ns: not significant, ****p < 0.0001.

### *Salicornia-*treated fruit flies are lighter and have reduced triacylglyceride levels

To assess whether feeding 0.2 % SEE affects the body composition of mated female flies, we quantified parameters such as body weight, triacylglyceride levels (TAG), protein, and glucose content after three weeks of treatment. Regarding body weight, the SEE-treated group was slightly lighter than the non-treated animals (Fig. 2A, p=0.01). We also quantified the TAG levels and found substantially lower TAG amounts (Fig. 2B, p = 0.007), while for the body protein content, the experimental groups did not show any significant differences (Fig. 2C). Moreover, we found no significant changes in glucose levels in SEE-treated animals (Fig. 2D). Finally, because SEE-treated flies had lower TAG levels, we analyzed the starvation resistance, which is usually directly associated with the body fat content^23^. In good agreement, SEE-treated animals showed a reduced starvation resistance with a median survival of 37 h compared to controls with a median survival of 41 h (p < 0.001; Fig. 2E). We next analyzed if the addition of the SEE changed the survival in response to desiccation stress with a similar median lifespan for the control group and flies after two weeks of treatment (Fig. 2F, p = 0.0633).

**Figure 2:**
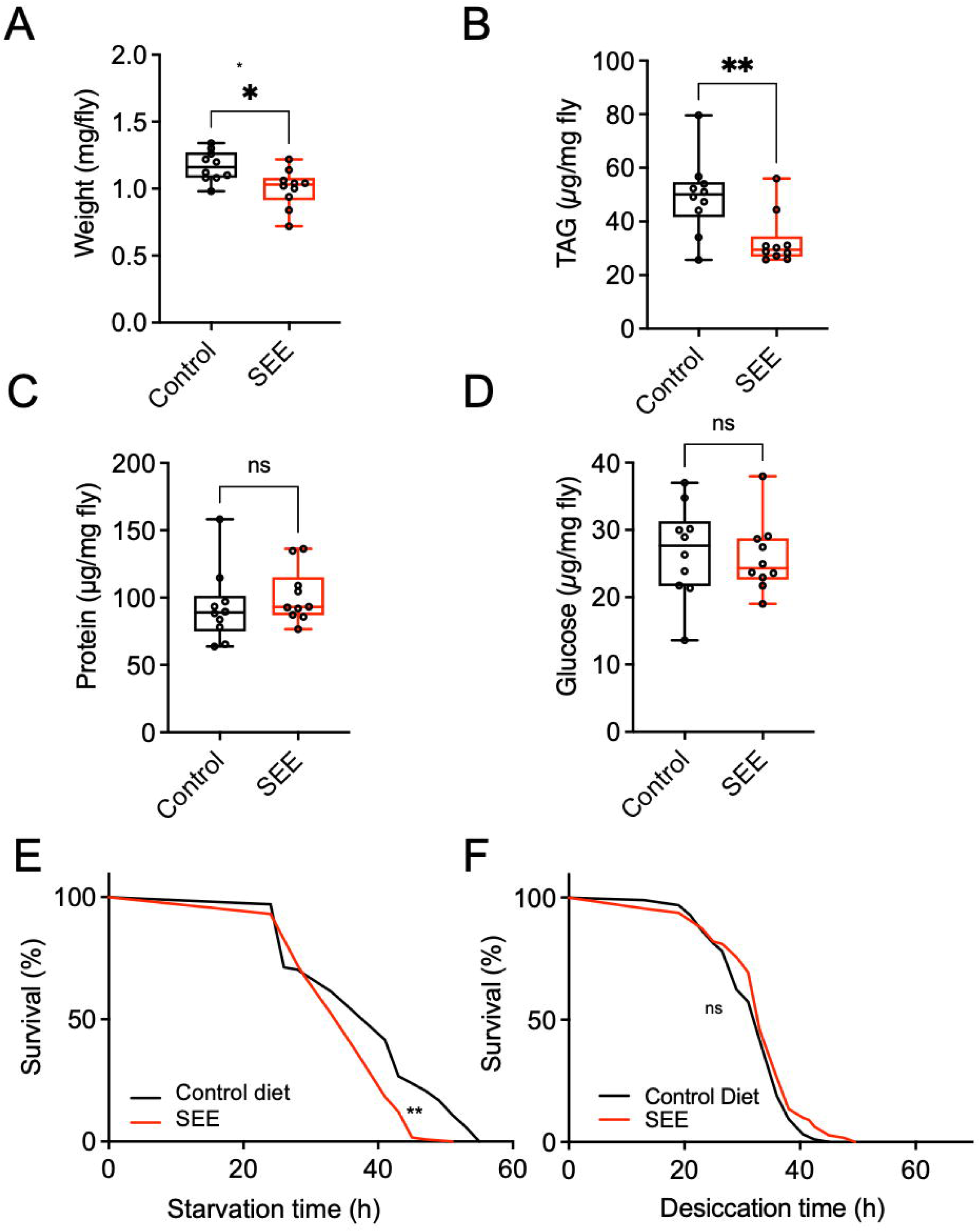
Body composition and resistance towards starvation and desiccation of female flies in response to SEE. (A) Body weight of female flies that were treated with SEE or not (N = 10, box plot showing median and minimum as well as maximal values). (B) Effects of SEE treatment on body fat (triacylglycerol) levels (N = 10). (C) Quantitative analysis of the protein content of SEE-treated females and their matching controls (N = 10). (D) SEE effects on the glucose content of flies (n = 10). E) Starvation resistance of flies treated for three weeks with SEE. (N > 100, median survival of SEE-treated flies = 37 h, median survival of untreated flies = 41 h). F) Desiccation resistance of SEE-treated female flies (N = 100, median survival of SEE-treated flies = 33 h, median survival of untreated flies = 33). SEE: *Salicornia europaea* extract, ns: not significant.

### Effect of SEE extract on energy metabolism

We quantified the nutritional intake using the blue food assay to exclude indirect effects caused by reduced uptake, thereby leading to an induced caloric restriction with its lifespan-prolonging effects. There were no statistically significant differences in food consumption for 24 hours between the experimental groups (Fig. 3A, p = 0.5681), which implies no change in energy intake. We also quantified the most energy-consuming trait in *Drosophila*, egg production, to evaluate if the observed lifespan extension compromised other health measures or came at the cost of reduced fecundity. The measurements revealed that the SEE-treated flies showed a higher reproductive output as evidenced by higher numbers of eggs laid in the two-week monitoring period (Fig. 3B, p = 0.0146). In addition, we analyzed the animals’ metabolic rate, and found that the experimental groups showed no significant alteration in CO_2_ production (Fig. 3C). Finally, we quantified physical activity as a measure of energy expenditure over 24-hour periods using the *Drosophila* activity monitor. Here, no significant differences were seen between the experimental groups (Fig. 3D).

**Figure 3:**
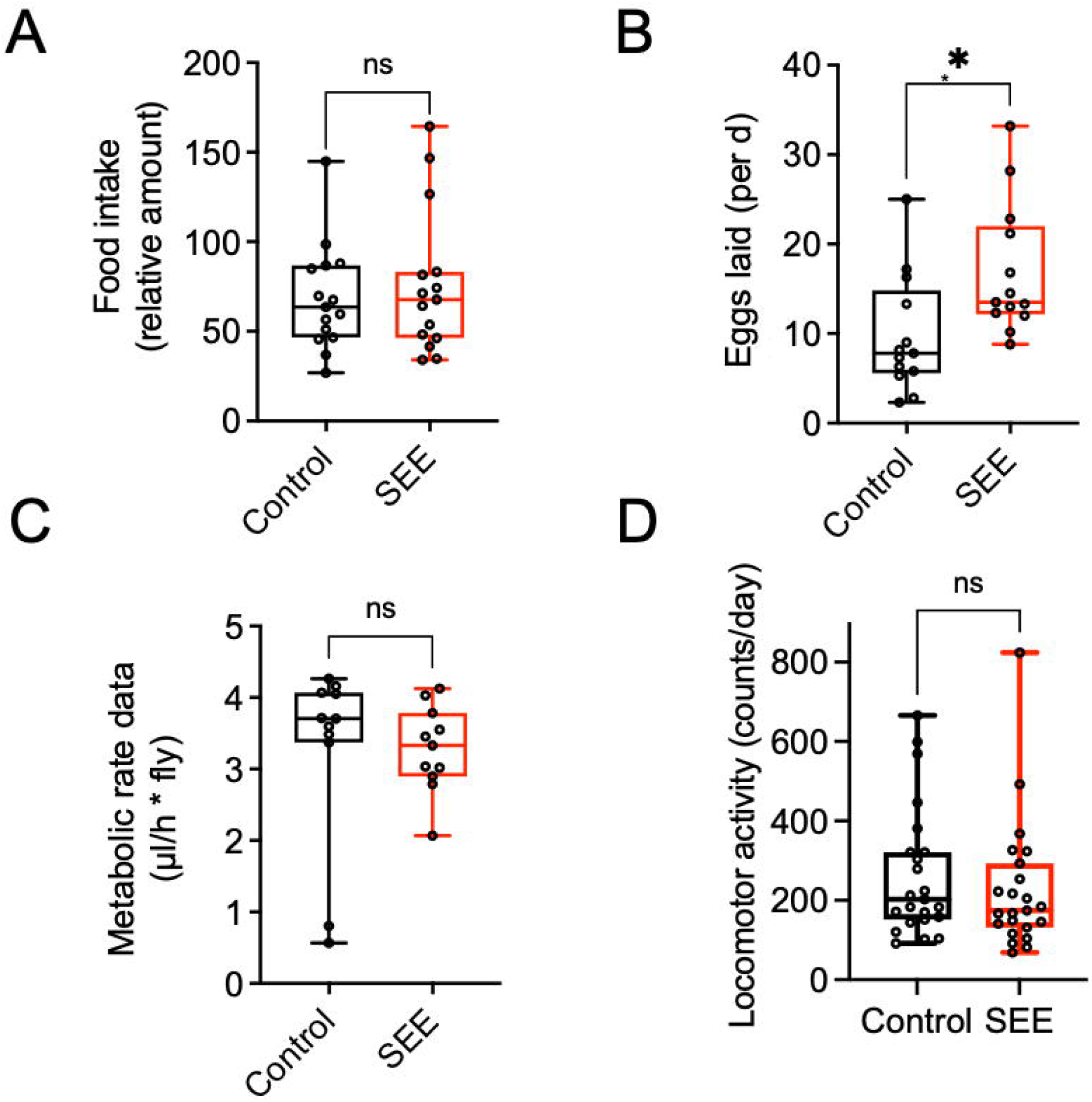
Energy intake and energy expenditure in response to SEE. (A) Food intake of female flies in response to SEE treatment during a 24 h period (N = 15). (B) Influence of SEE on egg production over a period of 14 days, shown as eggs laid per day (N = 10). C) Metabolic rate of flies subjected to SEE (N = 10). (D) The locomotor activity of adult flies either subjected to SEE treatment or not was quantified using a DAM monitor during 24 h periods (N = 25).

### Stress resistance to high-fat dieting is enhanced after SEE treatment

We next tested the interaction of SEE with two major nutritional stressors, namely high-fat and high-sugar diets. Flies treated with 0.2% SEE had the same median lifespan as the control group if confronted with a high-sugar diet (Fig. 4A, p = 0.7368). In contrast, when rearing flies on food containing 20% coconut oil (high-fat diet), we found that the SEE addition prolonged the median lifespan by about 16.6% from 30 d under control conditions to 35 d in response to SEE application (Fig. 4B; p < 0.0014). The maximum lifespan also increased significantly from 48 d in controls to 51.5 d in the SEE group (p = 0.0180). Interestingly, these results are in line with the extracts’ robust dose-dependent lipase inhibiting activity indicating a mode of action for the extract, which involves targeting the intestine lipases thus reducing intestinal absorption of dietary fat. When we employed an in vitro assay to examine the lipase-inhibiting properties of the SEE, a 1:2 dilution of SEE reduced lipase activity by almost 90%, and even a 1:20 dilution led to an enzyme inhibition of almost 20% (Fig. 4C).

**Figure 4:**
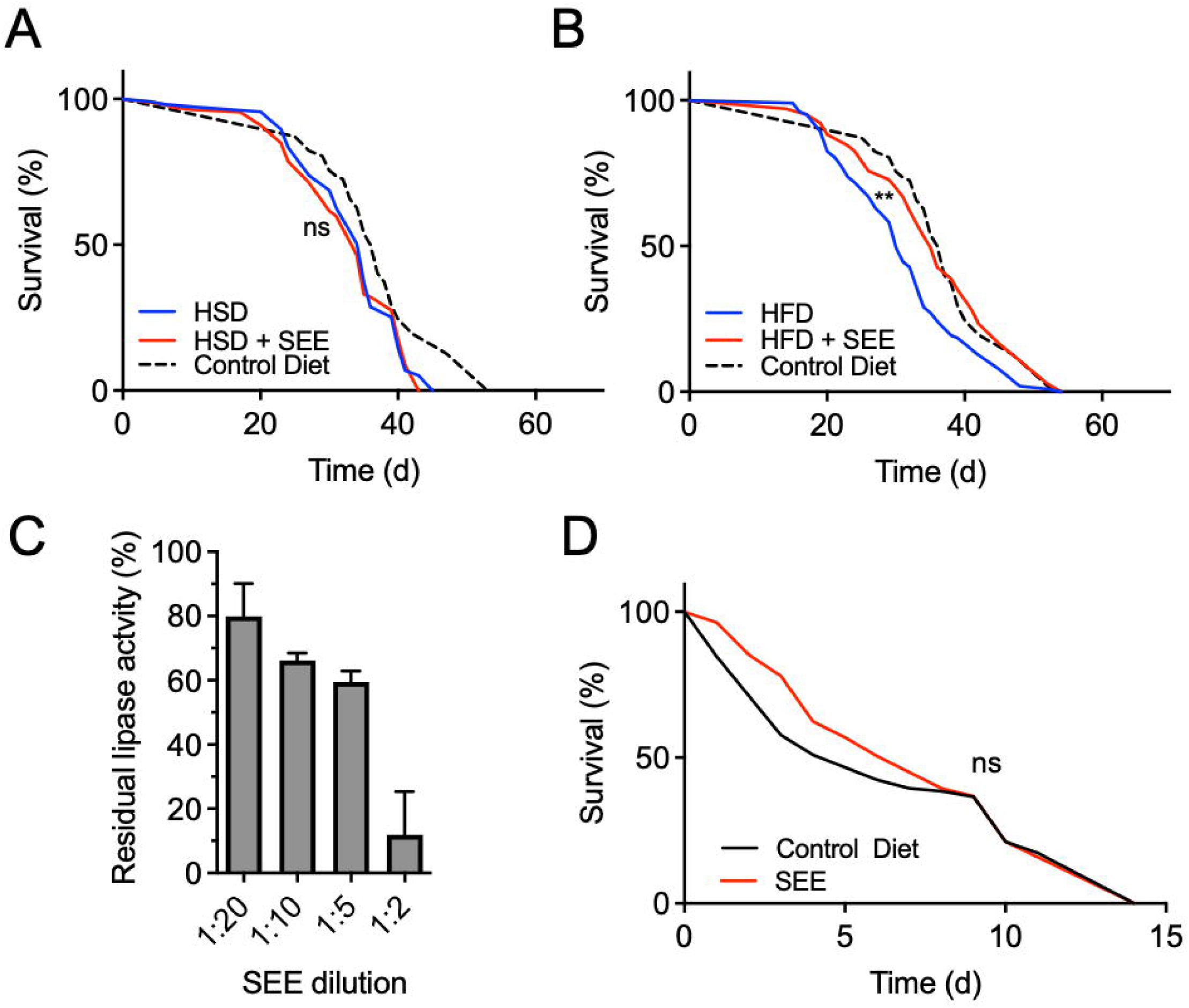
Stress resistance to high-fat dieting is enhanced after SEE treatment. (A) Impact of SEE on the survival of flies subjected to a high-sugar diet (N > 110, the median lifespan of SEE-treated = 34 d, the median lifespan of untreated flies = 35 d). (B) The survival of flies fed with SEE and subjected to a high-fat diet (N = 100, the median survival of SEE-treated flies under HFD = 35 d, the median survival of untreated flies under HFD = 30 d). (C) SEE extract affects lipase activity in a dose-dependent manner (N = 3). Listed are different SEE dilutions and their effect on lipase activity. (D) The median survival of flies under paraquat stress is not affected by pre-feeding a SEE-containing diet for 3 weeks (N > 100, the median survival for SEE-treated flies = 7 d, the median survival of control flies = 6 d. SEE: *Salicornia europaea* extract, ns: not significant, **p < 0.01.

We also examined the effect of the extract upon paraquat treatment as an oxidative stress inducer. In this condition, we did not observe a lifespan-extending impact of the extract (Fig. 4D).

### Mode of action of the lifespan prolonging effects of *Salicornia* extract

To elucidate potential mechanisms by which SEE extends the lifespan of female flies, we next focused on the major signaling pathways involved in the aging process, including the Sir2 signaling pathway, the Insulin/IGF signaling (IIS) pathway, and the Tor signaling pathway. Since activation of the Sir2 pathway has life-prolonging properties in *Drosophila*^24^, we employed *Sir2*-deficient flies and observed an almost identical lifespan on the control diet if compared with the genetic background (*w*^*1118*^). Moreover, SEE treatment prolonged the lifespan of the *Sir2*-mutants, indicating that *Sir2* is not necessary to transmit the effects of SEE (Fig. 5A). Furthermore, we analyzed the Tor signaling pathway by using the hypomorphic allele *TOR^k17004^*. As expected^25^, we observed an increased lifespan in flies with a *TOR^k17004^* background compared to the matching control (*yw*). However, SEE-treatment did not further positively affect lifespan in these flies, showing that Tor-signaling is necessary for mediating the lifespan prolongation in response to SEE (Fig. 5B). Since the Tor signaling and IIS are interwoven^17^, we also assessed the effect of SEE on the latter signaling pathway. To this end, we evaluated the role of the FoxO transcription factor as a proxy of IIS signaling. *dFoxo*-deficient flies showed a significantly reduced lifespan compared to their matching controls (Fig. 5C). Administration of SEE extended the lifespan of these animals, which indicates that FoxO signaling is not required to mediate the SEE effects on lifespan (Fig. 5C). This increase in lifespan is substantially smaller than in wildtype flies, implying that FoxO signaling might be involved to some extent in SEE-mediated lifespan prolongation.

**Figure 5:**
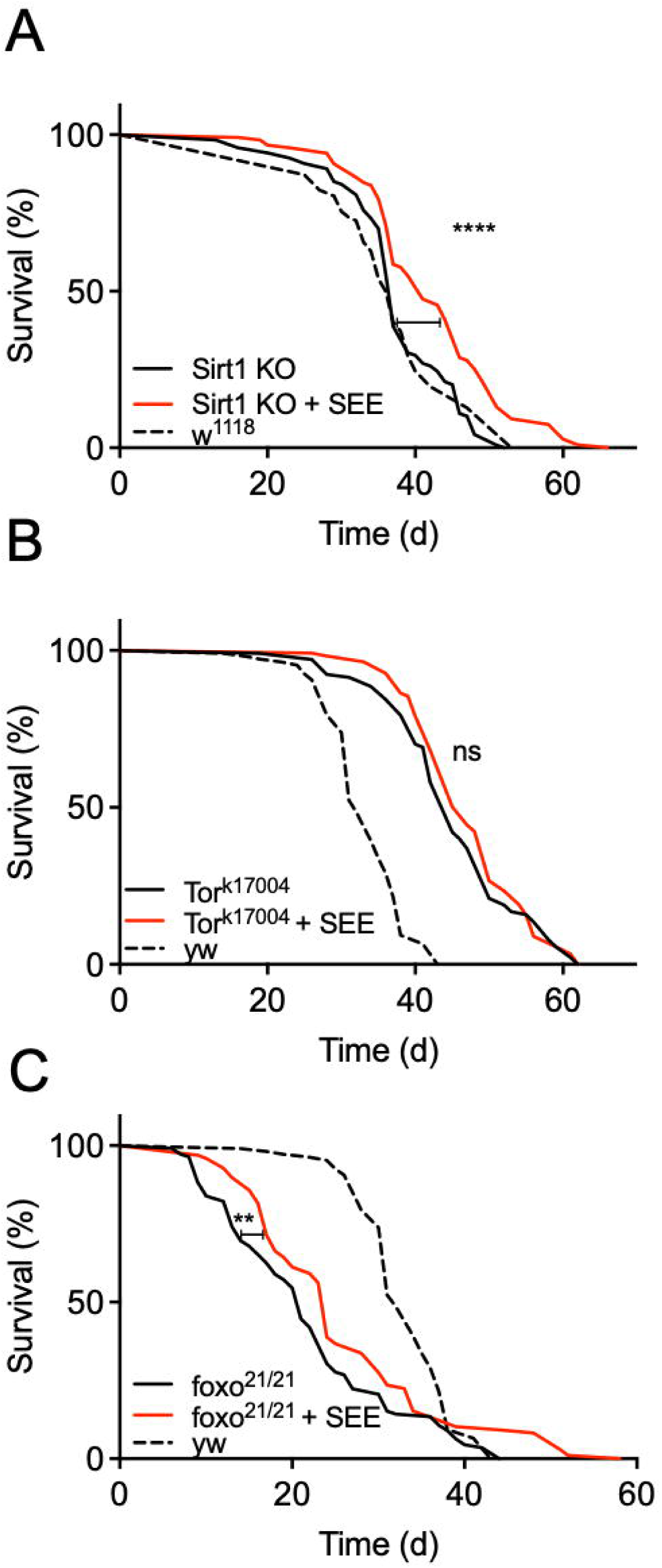
Signaling pathways potentially involved in mediating lifespan extension by SEE. (A) Lifespan of *Sirt1*-deficient flies in response to SEE in the diet compared with the untreated animals as well as with the genetic background controls (*w*^*1118*^) (N = 110, median lifespan of SEE-treated *Sirt1*-deficient flies = 41 d, median lifespan of *Sirt1*-deficient flies = 37 d). (B) Lifespan effects of SEE in *Tor* -deficient flies compared to non-treated flies as well as to the genetic background (*yw*) (N = 100, median lifespan of *Tor ^K17004^* flies = 45 d, median lifespan of *Tor ^K17004^* flies fed with SEE = 46 d). (C) SEE effects on lifespan of *dfoxo*-deficient animals compared with non-treated ones and with the genetic control (*yw*) (N = 100, median lifespan of SEE-treated foxo-deficient flies = 24 d, median lifespan of *dfoxo*-deficient flies = 21 d). **p < 0.01, ****p < 0.0001.

To further narrow down the mode and especially the side of action, we focused on the intestine and the fat body as the most important organs involved in mediating lifespan extension by nutritional interventions^17,26^. To test if the intestine is relevant for the lifespan extension mediated by SEE application, we performed a Smurf assay that is a direct proxy of intestinal health and correlates nicely with lifespan^27^. Here, at 20 days, 9.5% of the control group were smurf-positive, while the corresponding number for the SEE-treated group was only 1% (Fig. 6A). This difference was statistically significant (p < 0.005). We then manipulated Tor signaling in the enterocytes (*Np1-Gal4>UAS-Tor^DN^*) by induction of a dominant negative *Tor* allele (*Tor^DN^*). Here, SEE supplementation led to an extended lifespan of female flies (Fig. 6B). A similar result, i.e. a SEE-mediated lifespan extension, was obtained when we tested flies, in which Tor signaling was diminished through RNAi-mediated depletion of downstream S6K in enterocytes (*Np1-Gal4>UAS-S6K-RNAi)* (Fig. 6C). Furthermore, we used a driver specifically addressing intestinal stem cells and enteroblasts (*esg-Gal4*). Attenuation of Tor signaling using the Tor dominant negative allele in these cells did not abolish the beneficial effects of the extract (Fig. 6D). However, the lifespan prolongation was relatively small. Similarly, silencing of S6K in these cells (*esg-Gal4>UAS-S6K RNAi*) did not rescue the lifespan extension phenotype (Fig. 6E).

**Figure 6:**
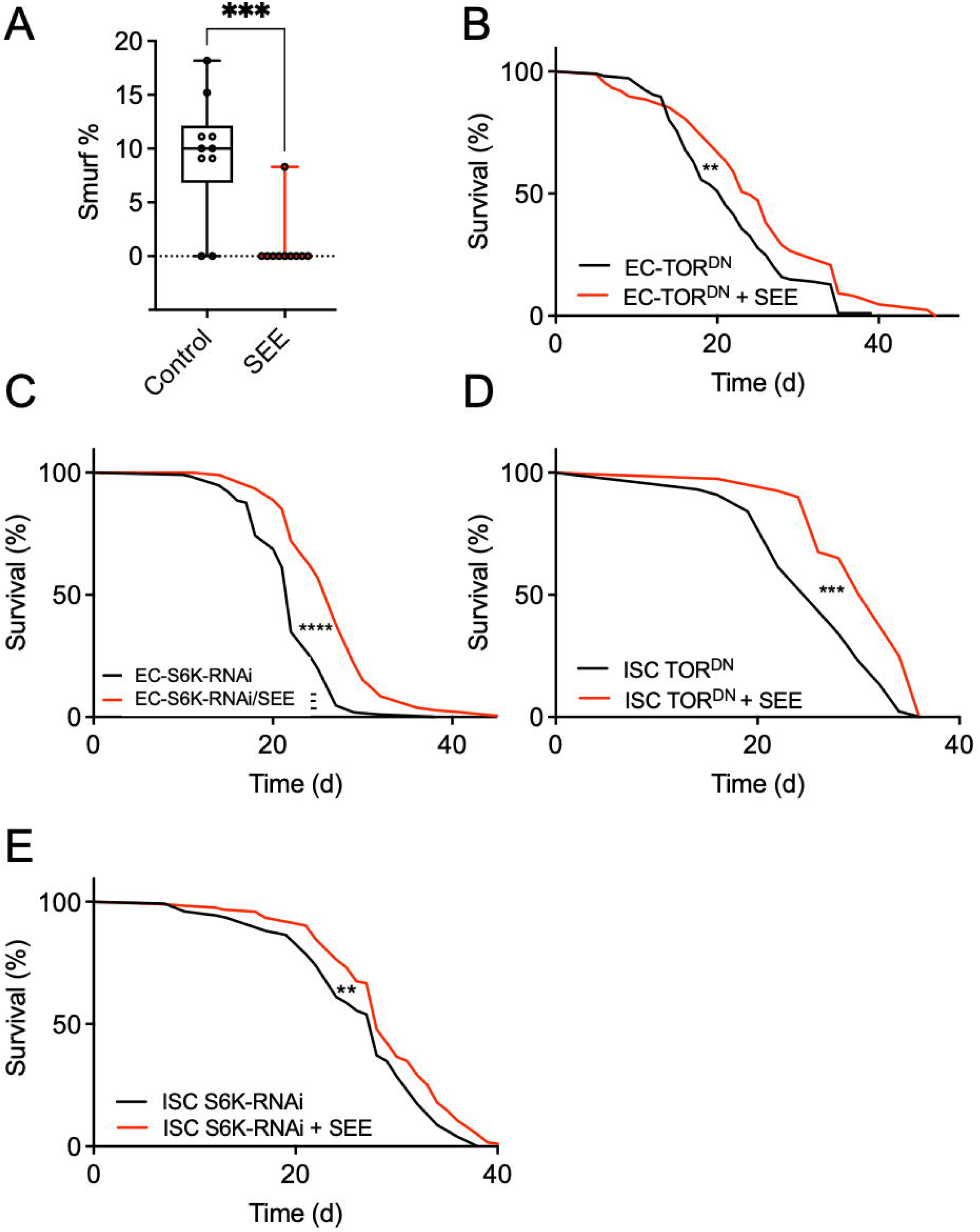
Tor/S6K signaling in different cell types of the intestine impacts SEE-induced lifespan effects. (A) Smurf analysis of flies fed a normal diet, or a diet supplemented with 0.2% SEE for 20 d. Data are shown as boxplots with whiskers indicating minimal and maximal values (N = 10). (B) The impact of SEE supplementation on the lifespan was examined in female fruit flies whose Tor signaling was reduced only in enterocytes (*Np1-Gal4/tuBGal80>UAS-Tor.TED*). Flies were maintained under control conditions or subjected to treatment with 0.2% SEE (N > 80). (C) The lifespans of fruit flies with reduced S6K expression in their enterocytes (*Np1-Gal4/tuBGal80>UAS-S6K-RNAi*) were determined in the presence and absence of 0.2% SEE (N > 100). (D) The SEE effect on lifespan was tested in flies with dominant negative Tor expression (Tor^DN^) in ISCs and EBs (*esgGal4, UAS-GFP; tub-Gal8>UAS-Tor.TED*) (N = 45, median lifespan of *esgGal4, UAS-GFP; tub-Gal80>UAS-Tor.TED* + SEE = 31 d, *esgGal4, UAS-GFP; tub-Gal80>UAS-Tor.TED* = 26 d) and (E) in flies with RNAi-depleted S6K in ISCs and EBs (N > 120, the median lifespan of *esgGal4, UAS-GFP; tub-Gal80>UAS-S6K-RNAi* + SEE= 28, the median lifespan of *esgGal4, UAS-GFP; tub-Gal80>UAS-S6K-RNAi* = 28). ns: not significant, *p < 0.05, **p < 0.01, ***p < 0.001, p < 0.0001.

To evaluate if Tor-signaling in the fat body is required for the SEE-induced lifespan extension, we used the RU486-inducible abdominal fat body-specific driver (*P (106) GS* ) for expression of the *Tor^DN^* allele (Fig. 7A). SEE application led to a significant increase in lifespan under these conditions, implying that Tor signaling in the fat body is not required for the SEE effect on lifespan (Fig. 7B). We also silenced S6K by RNAi targeted to the fat body of female flies. In good alignment, these animals showed prominent lifespan extension by SEE application (Fig. 7B) suggesting the mode of action of the extract was uncoupled from downregulation of S6K in the fat body.

**Figure 7:**
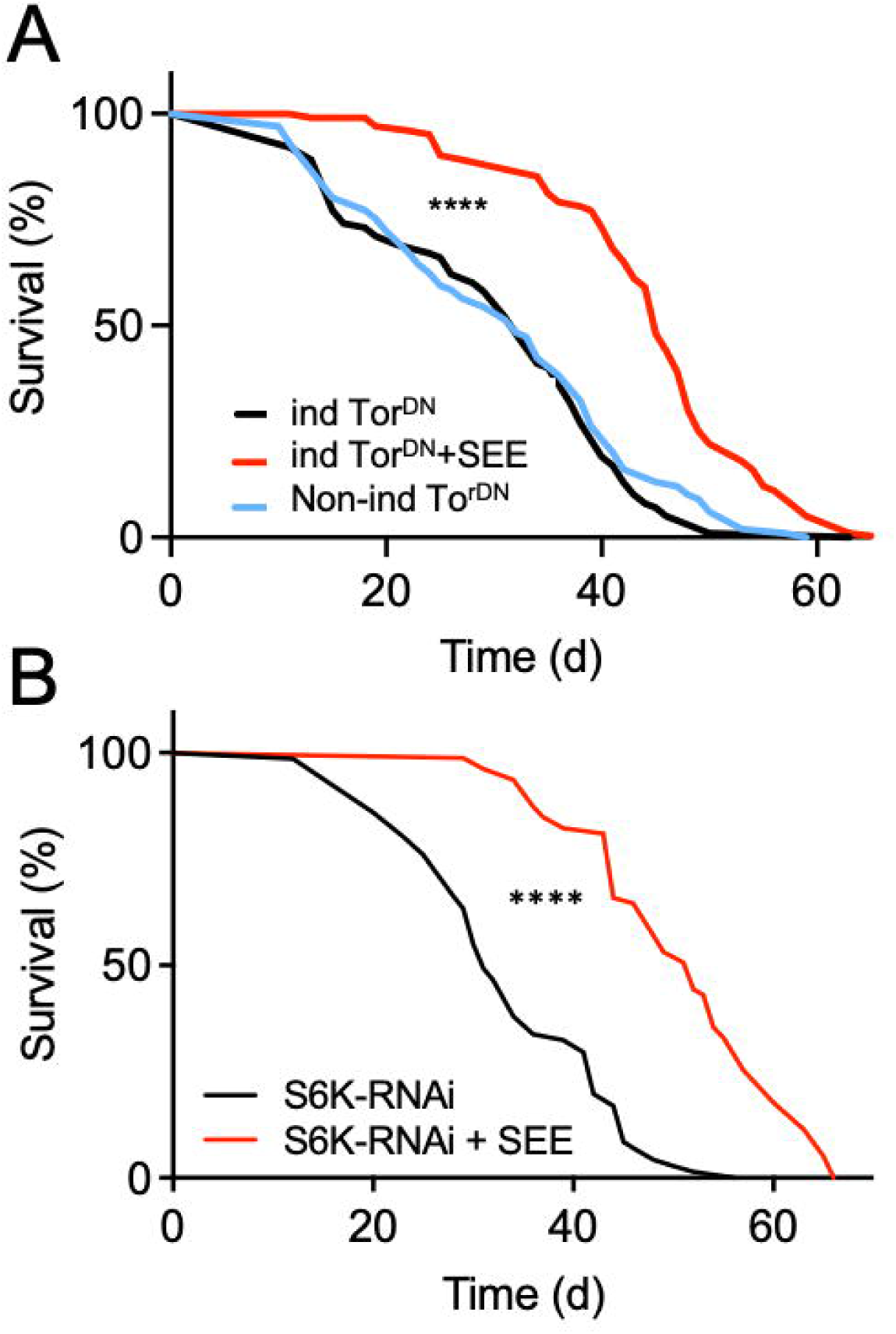
Fat body Tor/S6K signaling has no impact on lifespan prolongation induced by SEE. (A) Lifespan of flies with abolished Tor signaling (Tor^DN^; ind for induced) in the fat body (*P(GS)106>UAS-Tor.TED*) with and without SEE treatment, and of non-induced flies of the same genotype (N = 100, median lifespan of uninduced flies = 32 d, median lifespan of induced flies = 32 d). (B) SEE effects on lifespan of flies with reduced S6K signaling in the fat body compared with non-treated animals (*ppl-Gal4>UAS-S6K-RNAi*) (N > 70, median lifespan of *ppl-Gal4>UAS-S6K-RNAi* = 31 d, median lifespan of *ppl-Gal4>UAS-S6K-RNAi* + SEE = 52 d).

## Discussion

We found that an aqueous extract of the marsh samphire (*S. europaea*) (SEE) significantly prolongs the lifespan of female *D. melanogaster*. This significant lifespan extension by 30-40% is seen for mean and maximum lifespan. Since we observed this effect in more than one *Drosophila* strain, the extract possibly exhibits similar positive effects on fruit flies in general. Only females show this SEE-mediated effect on lifespan, whereas male flies did not benefit from a SEE treatment. Comparable sex differences in response to a life-prolonging intervention have already been shown in other studies^28,29^. Here, mainly mice and *Drosophila* served as models. Still, a general picture emerged that females benefit more from lifespan-extending interventions than males, regardless of whether they are pharmacological or nutritional^28–30^. In a recent study focusing on the extract of the brown alga *E. bicyclis* , we already showed a sex-specific lifespan extension in response to this nutritional/pharmacological intervention^17^. This clear dependence differs, for example, from a recent study with extracts from a brown alga *S. polyschides*, in which both sexes showed similar degrees of lifespan extension^16^. Despite this wealth of studies showing sex-specific effects on longevity, the underlying mechanism for these differences usually remained unclear. However, one of the few mechanistic studies attributed the differential impact of a life-prolonging pharmacological intervention to specific properties of the *Drosophila* intestine^26^. A similar mechanism is also operative in response to the SEE extract, as our data imply that the lifespan-prolonging effects depend on modifications within the gut.

We also elucidated the underlying molecular mechanism responsible for SEE-mediated lifespan. In the process, it turned out that the Tor signaling pathway is indispensable for these very effects. This finding is in line with several studies in which the importance of the Tor signaling pathway for all age-associated processes has been worked out^31–33^. Consequently, targeting this signaling pathway to enhance life- and health-span is a promising strategy. In further experiments, we tried to identify the organ responsible for the lifespan extension induced by SEE. The intestine and the fat body were the prime candidates for this role. Our smurf experiments showed that SEE treatment positively affects intestinal functionality. This reaction is relevant as barrier loss in the intestine is a hallmark of aging and is closely associated with premature death^27,34^. Silencing Tor signaling in different gut compartments showed that the SEE-induced response was not completely abolished. Still, it was substantially reduced in animals experiencing blockade of Tor-signaling in intestinal stem cells. This dependency implies that the intestine is involved in SEE-induced lifespan prolongation, but it is not the only target organ. The fat body has often been shown to be relevant for lifespan-extending interventions^24^. Nevertheless, it is dispensable for mediating the SEE-induced effects on lifespan. Moreover, we found a lipase-inhibiting activity of SEE, which is presumably not the primary reason for the lifespan prolongation under control conditions but might have an additional beneficial effect, especially in high-fat dieting situations.

If we compare the main results of the current study using a *Salicornia* extract with those of a previous one employing an *E. bicyclis* extract^17^, several commonalities are striking. The effects observed in both sets of experiments comprise strict sex-specificity and dependence on Tor signaling. The substantially reduced life-prolonging properties in a *dfoxo*-deficient background imply that FoxO signaling is important but not essential for mediating the SEE effects. This differs from the *Eisenia* extract, where the lifespan prolongation is strictly FoxO-dependent. Furthermore, the different extracts induce increased resistance to different nutritive stress situations. While the *Eisenia* extract protected particularly well against a high-sugar diet and hardly affected a high-fat diet, the *Salicornia* extract was particularly effective in mediating positive effects when the animals were confronted with a high-fat diet. This result implies that the molecular mechanisms underlying the effects of *E. bicyclis* and *S. europae*extracts share the requirement of active Tor signaling but differ in how they interfere with this signaling pathway.

*Salicornia* species are potential food sources and a source for pharmaceutically active compounds^19^. An essential reason for using *Salicornia* in this way is the almost worldwide availability of this plant. As a source for pharmaceutically relevant phytochemicals, *Salicornia* species have been studied intensively. The abundance of active compounds in the extract might account for the unexpected increased fecundity of SEE-treated female flies despite reduced fat content and increased lifespan which is reminiscent of the pro-longevity and - fecundity effects of the TCA intermediary metabolite citrate. In line with this assumption, SEE is enriched with two TCA metabolites, namely, isocitrate and fumaric acid^35^.

Highly relevant for this work are the metabolic effects of *Salicornia* products, with particular emphasis on antidiabetic properties and those leading to a reduction in fat storage^36^. Lowering fat deposition was associated with interference with SREBP1-mediated processes induced by different components found in the *Salicornia* extract. This reduction in fat storage is entirely in line with our observations, showing a potent inhibition of lipase activity. Moreover, immunomodulatory and anti-inflammatory effects were identified, predominantly mediated by polysaccharides of *Salicornia* extracts^37^.

The anti-obesogenic activity of the SEE is a highly desirable feature. The search for plant extracts and other sources of compounds with precisely these properties has high priority^38^. Here, different mechanisms are operative^39^. Lipase inhibition, as observed in the SEE, appears to be a significant mechanism of anti-obesogenic effects induced by natural products^40^. Despite this compelling bioactivity of the SEE as an anti-obesogenic source, the lifespan-prolonging effects are operative through a different mechanism, which is Tor-dependent. The control diet, where SEE induced a substantial lifespan prolongation is not rich in lipids (hier vielleicht einen Wert nennen, siehe angehängte Datei), further pointing to different mechanisms underlying the lifespan prolongation and the anti-obesogenic effects.

Metabolomic analysis including SEE^41^ revealed several different components that have exciting properties. Here, we compared the metabolomic profiles of aqueous extracts from two brown algae (*S. polyschides* and *E. bicyclis* ) with the SEE. All of them could increase lifespan in a *Drosophila* model, but they induce this effect through different mechanisms. Therefore, we will discuss especially those metabolites that we found explicitly in the *Salicornia* extracts and are known to affect lifespan. Chlorogenic acid, tuberonic acid, melleolide, isorhamnetin, or kaempferol are especially relevant here. Chlorogenic acid, for example, is one of these substances that showed an apparent life-prolonging effect in *C. elegans*. Here, interaction with insulin signaling and the Akt-FoxO axis was required to unfold these effects ^42,43^. In addition, isohamnetin, a methylation product of quercetin, also enhances lifespan in *C. elegans* together with resistance to stressors^44^. A Lotus (*Nelumbo nucifera*) stamen extract containing several of these compounds, found explicitly in the *Salicornia* extract, showed a robust delay of aging in yeast. Here, especially kaempferol and isohamnetin must be mentioned ^45,46^. Thus, some of the lifespan-prolonging effects of the *Salicornia* extract could be mediated through the compounds discussed above that are present at high concentrations in the SEE. We intend to use this information, which ultimately paves the way for identifying the lifespan-mediating metabolites, for this very purpose in follow-up studies applying purified test compounds alone and in combination. This result is beyond the scope of this manuscript and should be the subject of future studies.

In conclusion, the aqueous *Salicornia* extract we used in this study shows a vast, sex-specific lifespan extension in *Drosophila*. On the one hand, this opens a broad field of application for use as a human nutritional supplement. On the other hand, mechanistic studies can provide information about the mode of action of the life extension and thus provide a crucial first step toward identifying the life-extending substances of the extract. The molecular basis of the sex-specific effect of life extension can also be elucidated in the future.

## Methods

### Plant extraction

*Salicornia* extract was prepared according to the method described by Onur et al.^11^. In short, the dried algal material was ground with an analytical mill (IKA Type A11 basic). Three grams of ground alga were transferred into a test tube containing 30 ml of boiling double distilled water. After slight stirring, the suspension was sonicated for 1 min and centrifuged for 2 min at 2000 x *g*. The supernatant was filtered, and the soluble extract was freeze-dried and stored frozen until use.

### Fly husbandry

Wild-type adult flies were kept as previously described^17,47^. In brief, they were cultivated on a diet containing 5% (w/v) yeast extract, 8.6% (w/v) corn meal, 5% (w/v) sucrose, and 1% (w/v) agar-agar supplemented with 1% (v/v) propionic acid and 3% (v/v) nipagin. Adults, 3–5-days after hatching were used in the experiments. All experiments were performed at 25 °C, a light dark cycle of 12 h: 12 h, and 60% humidity. For most experiments, the wild type strain *w*^*1118*^ was used. The following fly strains were used for the study: *w*^*1118*^ Bloomington stock ID #5905), Sir2-deficient (Bloomington stock ID #8838), Tor deficient (Bloomington stock ID #11218), dfoxo-deficient (foxo^21^*^/^*^21^) (Marc Tatar lab), *y^1^w*^*1118*^ (Bloomington stock ID #111281), *P (Switch)106* (Ronald Kühnlein), *NP1-Gal4* (D. Ferrandon, Straßburg), *esg-Gal4*, *ppl-Gal4* (P. Leopold), *UAS-Tor.TED* (Bloomington stock ID #7013), *UAS-S6K-RNAi* (Bloomington stock ID #41702).

### Lifespan assay and analysis

Lifespan assays were essentially performed as described earlier^48^. The mated female flies were separated into experimental groups and kept on the food described above. Vials were regularly changed every 2 days and dead flies were counted every day. For the high-sugar diet, the sucrose concentration was increased to 30%, and for the high-fat diet; the concentrated medium was supplemented with 20% palm fat. To supplement the food with the algal extract, the aqueous extract was added on top of the food in the experimental vials at the final concentration of interest. Then, the water is allowed to evaporate to dry the food. At least 100 animals were used at a minimum per experiment.

### Body fat quantification

The whole-body triacylglycerol (TAG) content was measured using coupled colorimetric assay as previously described^23,48^. In short, samples were collected (5 females per sample) and weighted. 1 ml PBS/Tween-20 (0.05%) was added to the samples, and they were homogenized in a bead ruptor apparatus (BioLab Products, Bebensee, Germany) for 2 minutes at 3.25 m/s. Next, samples were centrifuged for 3 minutes at 3000 x *g* and the supernatant was transferred to new tubes. The supernatant was heat-inactivated at 70 °C for 10 minutes and centrifuged for 3 minutes at 2500 x *g*. 50 μl of each sample were added to a 96-well plate and the absorbance was measured at 500 nm (T_0_). 200 μl prewarmed triglyceride reagent were added to each well and incubated for 30 minutes at 37 °C with mild shaking. The absorbance was again measured at 500 nm (T_1_). The TAG concentration was determined by subtracting T_0_ from T_1_. The TAG content was quantified using a triolein-equivalent standard curve.

### Glucose measurement and protein content analysis

The body glucose levels were determined using a Glucose (HK) Assay Kit (GAHK-20, Sigma-Aldrich, Taufkirchen, Germany) according to the manufacturer’s instructions with minor modifications. Samples were collected (5 female flies per sample), weighted, and homogenized using a bead ruptor apparatus (BioLab Products, Bebensee, Germany) for 2 minutes at 3.25 m/s. For the glucose measurement, the supernatants were heated for 10 minutes at 70 °C and then centrifuged for 3 minutes at 4 °C. 30 μl of the supernatant were added to a well of a 96-well plate. 100 μl of the HK reagent were added to each well and the plate was incubated at room temperature for 15 minutes. Then, the absorbance was measured at 340 nm. The glucose content was calculated using a glucose standard curve. For the determination of the protein content, the samples were centrifuged for 1 min at 1000 rpm(x *g*) at 4C. The supernatant was transferred to a new 1.5 ml tube and centrifuged again at 6000 rpm(x *g*), at 4 °C for 10 minutes. The supernatant was again transferred to new tubes and centrifuged at maximum speed, at 4 °C for 10 minutes. The protein content was measured using the Pierce BCA Protein Assay Kit according to the manufacturer’s instructions.

### Starvation, desiccation, paraquat resistance, and fecundity assays

In order to assess flies’ resistance to starvation, flies were transferred to vials containing 1% agar after feeding with food of interest. The survivorship of the flies, which were kept at 25 °C, 60% humidity, and 12/12 hours light/dark cycle, was recorded every two hours. To determine the resistance to desiccation, flies were transferred to empty vials after feeding them with the food of interest. Again, the flies were kept at 25 °C and 60% humidity and 12/12 hours light/dark cycle. The survival of flies was monitored every 1 to 2 hours.

For the paraquat assay, the flies were treated with the extract for 3 weeks and then transferred to food supplemented with 20 mM paraquat (methylviologen) without extract addition. The number of dead flies was counted for 2 weeks.

To determine egg production, mated female flies were kept in vials containing the food of interest and the laid eggs were counted daily for 14 days.

### Food intake

The food consumption of flies was measured using the consumption-excretion method^49^. The fly medium or blue dyed medium (0.5% (w/v) Brilliant Blue FCF food dye; E133) was dispensed into caps of 2 ml screw cap vials. For adaptation, individual flies were transferred to 2 ml screw cap vials with CM. After a few hours feeding on the concentrated medium, the flies were transferred to 2 ml vials with blue dyed food. After 24 hours feeding on blue dyed food 3 ceramic beads and 500 μl H_2_O were added to the vials. The samples were homogenized using a bead ruptor (OMNI International, OMNI Bead Ruptor 24) for 90 sec at 3.25 m/s. The homogenized samples were centrifuged at 3000 rpm(x *g*) for 3 minutes to deposit the tissue debris. Then 200 µl of the supernatant was added to clear 96-well plates and the absorbance was measured at 630 nm.

### Smurf assay

Animals were reared on fly medium? for 20 and 27 days, before they were transferred to a dyed medium containing 1.5 % (w/v) Brilliant Blue FCF in CM. After 24 hours, the animals were inspected and counted as Smurf when the blue dye was observed outside of the intestine ^27^.

### Statistics

Statistical analyses and plotting figures were done by GraphPad Prism versions 7 and 8. For lifespan analyses a Log-rank (Mantel-Cox) test was used. For other experiments, first the data were tested for normal Gaussian distribution using the Shapiro Wilk normality test. Subsequently, unpaired t-test was used for data exhibiting normal distribution, and the Mann-Whitney test was used for the other data.

## Funding

This work was supported by the German Research Foundation as part of the CRC1182 (project C2) and by the BMBF (Project “Vision Epifood”).

## Acknowledgments

The marsh samphire extract was developed based on “The Plant Extract Collection Kiel in Schleswig Holstein – an open access screening Library” kindly provided by Prof. Frank Döring (Kiel, Germany, Coordinator of the BMBF project “Vision-Epifood”). We would like to thank Britta Laubenstein and Christiane Sandberg for excellent technical assistance and Dominique Ferrandon, Ronald Kühnlein, and Marc Tatar for flies.

## Conflict of interest

The authors declare that they have no conflict of interest.

## Notes

### Competing Interest Statement

The authors have declared no competing interest.

